# ASYMMETRIC EXPRESSION OF ARGONAUTES IN ARABIDOPSIS REPRODUCTIVE TISSUES

**DOI:** 10.1101/2020.05.18.102863

**Authors:** PE Jullien, DMV Bonnet, N Pumplin, JA Schröder, O Voinnet

**Affiliations:** Institute of Plant Sciences, University of Bern, Bern, Switzerland; Institute of Molecular Plant Biology, Swiss Federal Institute of Technology Zurich (ETH-Zurich), Zurich, Switzerland

**Keywords:** Argonautes, small RNA, reproduction, Gametes, endosperm, embryo

## Abstract

During sexual reproduction, development of a totipotent zygote from the fusion of highly differentiated gametes is accompanied by dynamic regulation of gene expression. This notably involves RNA silencing operated by Argonautes (AGO) effector proteins. While AGOs’ roles during *Arabidopsis* somatic life have been extensively investigated, less is known about their expression during reproduction, which proceeds via double-fertilization of an egg and a central cell, leading respectively to the embryo and a supportive tissue known as endosperm. Using full-locus translational reporters for all ten *Arabidopsis* AGOs, we uncover cell-specific expression patterns and AGO-intrinsic subcellular localizations in reproductive tissues. However, while some *Arabidopsis* AGOs are enriched in both male and female gametes, *i*.*e*. sperm and egg cells, they are comparably low-expressed in accessory, *i*.*e*. vegetative and central cells. Likewise, following fertilization, several AGOs are expressed in the early embryo, yet below detection in the early endosperm. Thus, there is pre- and post-fertilization asymmetry between the embryo and endosperm lineages. Later during embryo development, AGO9, AGO5 and AGO7 are restricted to the apical embryonic meristem in contrast to AGO1, AGO4, AGO6 and AGO10. Beside shedding light onto potential roles for RNA silencing during *Arabidopsis* reproduction, the plant material generated here should constitute a valuable asset enabling functional AGOs studies in many tissues beyond those involved in reproduction.

**Summary statement:** Arabidopsis genome encodes ten Argonautes proteins showing distinct expression pattern as well as intracellular localisation during sexual reproduction.

## Introduction

In most eukaryotes, sexual reproduction involves specialized cellular structures and entails complex orchestration of the timing and spatial localization of gene expression. In the model plant *Arabidopsis*, reproduction involves the development of a haploid structure, called gametophyte, in both male and female reproductive organs. The mature male gametophyte, or pollen grain, contains three cells: one vegetative cell and two gametes known as sperm cells. The mature female gametophyte, on the other hand, contains seven cells: three antipodal cells of unknown function, two synergides involved in pollen tube reception and guidance, one central cell and one egg cell. Fertilization of the egg cell by one of the sperm cells forms the embryo proper that later develops into the next-generation plant. Fertilization of the homodiploid central cell by the second sperm cell forms the endosperm, a nourishing tissue required for proper seed development. In addition, the female gametophyte as well as the developing endosperm and embryo are surrounded by 4 layers of maternal integuments connected to the mother plant’s vasculature at the chalazal pole.

Small RNAs (sRNA) are key regulators of gene expression. Their importance during sexual reproduction has been highlighted by the discovery of a class of reproduction-specific sRNA known as Piwi-interacting RNAs (piRNAs) in animals (Castel and Martienssen, 2013; Weick and Miska, 2014). Likewise, the importance of reproduction-specific small RNAs has also been recognized in plants (Mosher and Melnyk, 2010; Van Ex et al., 2011). Beyond the roles of microRNAs (miRNAs) in embryonic development, small interfering (si)RNA tame transposon in the pollen (Slotkin et al., 2009) and during ovule development (Olmedo-Monfil et al., 2010). siRNAs also might be linked to hybrid seed lethality as suggested by their implication in regulating parental genome dosage (Borges et al., 2018; Martinez et al., 2018). RNA silencing in plants can be divided into Post-Transcriptional Gene Silencing (PTGS) and Transcriptional Gene Silencing (TGS) processes (Bologna and Voinnet, 2014; Borges and Martienssen, 2015; Mallory and Vaucheret, 2010; Schröder and Jullien, 2019). Both pathways rely on the generation of small (s)RNAs of 21, 22 or 24 nucleotides in length by DICER-LIKE enzymes, of which there are four paralogs in *Arabidopsis* (DCL1-4). These sRNA execute PTGS or TGS upon their loading into ARGONAUTE (AGO) effector proteins. Despite the established impact of sRNA-pathway mutations in plant biology, little is known of the expression profiles of silencing-pathway proteins in reproductive and post-reproductive, early embryonic tissues. In particular, the cell-specific expression and subcellular localization patterns of AGOs remain largely unknown.

The *Arabidopsis* genome encodes ten AGO genes divided into three phylogenetic clades (Mallory and Vaucheret, 2010) conserved among flowering plants (Fang and Qi, 2015; Zhang et al., 2015). Clade I consist of AGO1/5/10, with AGO1 being the ubiquitously-expressed member. *ago1* mutants display aberrant sporophytic phenotypes likely due to AGO1’s key role in executing miRNA functions. Embryonic *ago1* mutant phenotypes were only observed, however, from the torpedo stage (Lynn et al., 1999). The second clade I member, AGO10, has been implicated in the control of shoot apical meristem (SAM) identity from late stages of embryo development (Lynn et al., 1999; Moussian et al., 1998), despite being expressed from very early stages (Tucker et al., 2008). In fact, early embryo phenotypes were only observed in the double *ago1ago10* mutant, thereby suggesting functional redundancy between the two proteins at this stage, possibly in executing miRNA-directed silencing. Supporting this notion, early embryo phenotypes are observed in *dcl1* mutants exhibiting compromised miRNA processing (Nodine and Bartel, 2010; Seefried et al., 2014; Willmann et al., 2011). Contrasting with that of AGO1 and AGO10, AGO5 expression is substantially enriched in reproductive tissues (Tucker et al., 2012). A dominant *AGO5* allele (*ago5-4*) arrests female gametophyte development altogether (Tucker et al., 2012) while *ago5* recessive mutants display early flowering (Roussin-Léveillée et al., 2020).

The AGO Clade II is constituted of AGO7, AGO2 and AGO3. AGO7 has a well-established role in leaf development, yet no evident function in reproductive tissues has been described thus far. Despite AGO2 and AGO3 displaying higher expression levels in reproductive tissue, their role(s) during reproduction remains, likewise, elusive because no overt developmental phenotypes were observed in the corresponding mutants (Jullien et al., 2020). Clade III includes AGO4/6/9/8, which, with the exception of the truncated AGO8 protein (Takeda et al., 2008), have all been implicated in DNA methylation commonly referred as the RNA-dependent-DNA-methylation (RdDM; (Matzke and Mosher, 2014). The RdDM pathway is required for the establishment of new DNA methylation pattern as well as the maintenance of cytosine methylation in the CHH context. AGO4 and AGO6 are mostly ubiquitously expressed, whereas AGO8 and AGO9 seem specific for reproductive tissue (Havecker et al., 2010; Olmedo-Monfil et al., 2010; Wuest et al., 2010). In particular, AGO9 is involved in meiosis, megaspore mother cell (MMC) differentiation and transposon silencing in the egg cell (Oliver et al., 2014; Olmedo-Monfil et al., 2010). MMC differentiation is also impeded in *ago4, ago6* and *ago8* mutants (Hernández-Lagana et al., 2016) suggesting an important role for RdDM during this process.

To help deciphering the roles of sRNA pathways in reproductive processes, we have generated stable transgenic lines expressing full length fluorescently tagged AGOs under their cognate endogenous promoter and analyzed their expression as well as intracellular localization in reproductive tissues.

## Results and Discussion

### Argonautes Translational reporters

In order to investigate the expression pattern and intracellular localization of the ten *Arabidopsis* AGO proteins, we generated full-locus reporter constructs in which the open reading frames (ORFs) of fluorescent proteins were fused to each AGO coding sequence in their genomic contexts, using multiple Gateway™ cloning. We engineered N-terminal translational fusions, since N-terminal, but not C-terminal, tagging preserves *Arabidopsis* AGOs’ functionality (Carbonell et al., 2012). Each AGO construct was cloned under the corresponding presumptive promoter (1.3kb to 2.5kb upstream start codons) and terminators (467bp to 1kb downstream of stop codons). This generated pAGO:FP-AGO constructs, where “FP” corresponds to either the Green Fluorescent Protein (GFP) or mCherry (mCh). For the sake of simplification, the constructs will be referred to FP-AGOX (where X is the number of each AGO1-10) in the main text and FPX in the figures. For example, pAGO1:mCherry-AGO1 will be shortened to mCherry-AGO1 in the main text and to mCh1 in figures. A detail map of the constructs can be found in Fig. S1 and a summary of their expression pattern in Fig.S11. Functional complementation was validated for AGO1, AGO4, AGO5, AGO6 and AGO7 and can be found in Fig. S2, Fig. S3, Fig. S4, Fig. S5 and Fig. S6 respectively.

### AGO Expression patterns in the mature female gametophyte

To gain insights into AGO protein expression patterns in the mature female gametophyte, we analyzed fluorescent signals of each protein fusion, using confocal microscopy. Almost all AGOs accumulate in the mature female gametophyte (Fig. 1A-K) except GFP-AGO10 (Fig. 1D) and mCherry-AGO3 (Fig. 1J), which are only detected in the maternal integument. The clade I mCherry-AGO1 and mCherry-AGO5 reporters display similar accumulation patterns in the female gametophyte (Fig. 1B-C). They are mainly detected in the egg cell and, to a lower extent, in the central cell. mCherry-AGO1 and mCherry-AGO5 also accumulate in the maternal integument, with mCherry-AGO1 being detected in both the inner and outer integument, and mCherry-AGO5 solely in the former, as previously reported (Tucker et al., 2012). Both mCherry-AGO1 and mCherry-AGO5 signals are particularly pronounced in the nucellus at the chalazal pole of the ovule. Clade I, GFP-AGO10 mainly accumulates in the inner-integument of the ovule with a stronger signal at the chalazal seed coat and in vascular tissues of the funiculus (Fig. 1D, Fig. S7A-B).

**Fig. 1.**
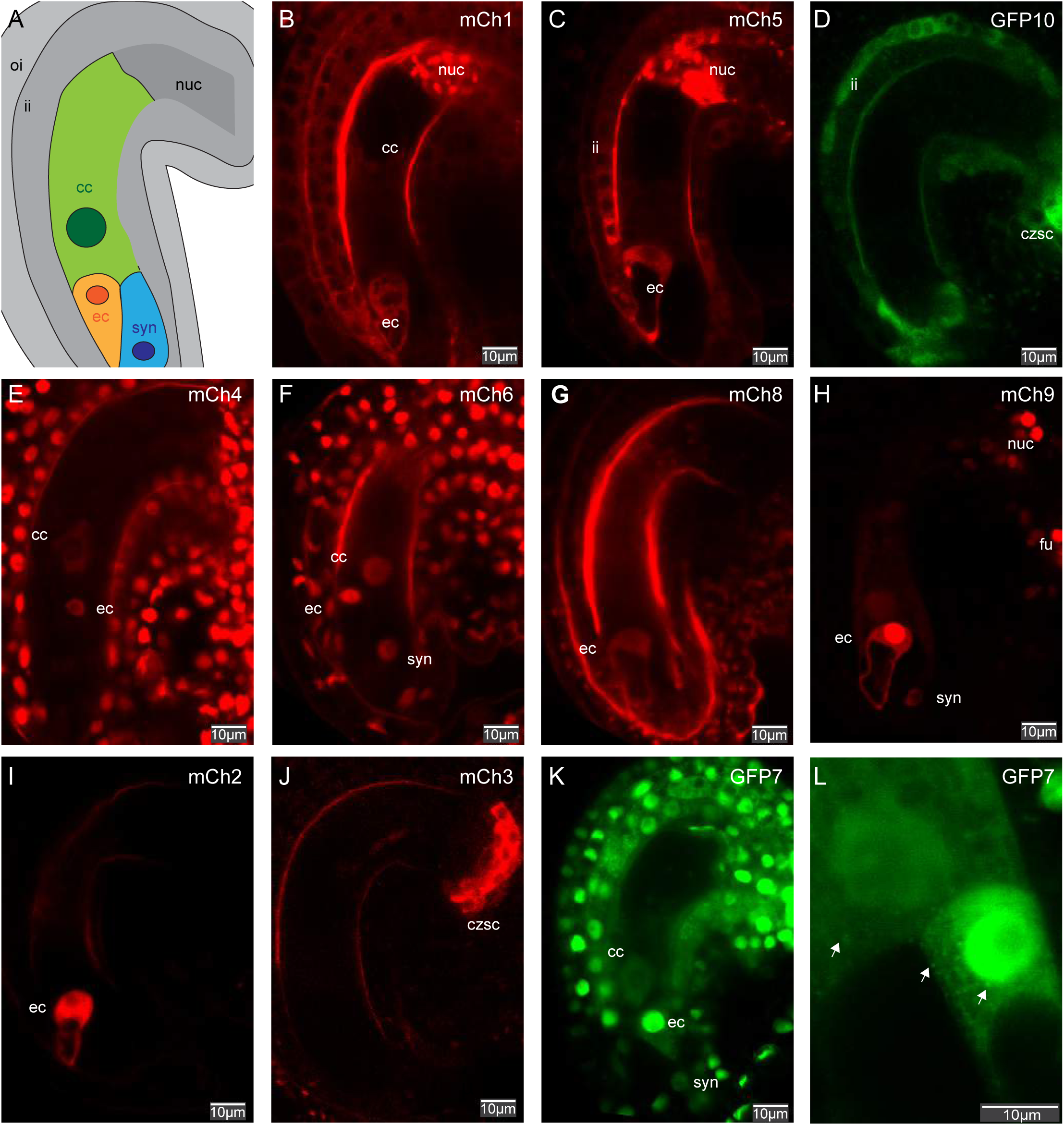
Argonaute expression in the mature female gametophyte. (A) schematic representation of a mature female gametophyte of Arabidopsis thaliana illustrating the three major cell type: the central cell (cc) in green, the egg cell (ec) in orange, the synergides (syn) in blue. The mature female gametophyte is surrounded by maternal sporophytic tissue represented in grey including the inner integument (ii), outer integument (oi) and the nucellus (nuc). (B-K) Confocal images representing the expression of the 10 Arabi-dopsis AGOs in mature female gametophyte: mCherry-AGO1 (B), mCherry-AGO5 (C), GFP-AGO10 (D), mCherry-AGO4 (E), mCherry-AGO6 (F), mCherry-AGO8 (G), mCherry-AGO9 (H), mCherry-AGO2 (I), mCher-ry-AGO3 (J) and GFP-AGO7 (K). (L) Confocal image illustrating the intra-cellular localization of GFP-AGO7 in the egg cell and central cell. Scale bars represent 10μm.

The main clade III *i.e*. RdDM AGOs, mCherry-AGO4 and mCherry-AGO6, accumulate ubiquitously in both integuments and female gametophyte, with stronger signals in the egg cell (Fig. 1E-F). The reproduction-stage-specific RdDM AGO’s, mCherry-AGO9 and mCherry-AGO8, show more spatially-restricted expression patterns (Fig. 1G-H). mCherry-AGO9 accumulates mainly in the egg cell within the female gametophyte but can be detected, albeit at lower levels, in the central cell. Strong mCherry-AGO9 accumulation is also detected in the nucellus and funiculus (Figure1H and Fig. S8A), consistent, for the latter, with previous *in situ* hybridization results (Olmedo-Monfil et al., 2010). mCherry-AGO8 is specifically detected in the mature egg cell (Fig. 1G). Egg cell expression of both AGO8 and AGO9 is supported by conclusions drawn from transcriptional fusions for both AGO9 and AGO8 (Sprunck et al., 2019) as well as transcriptomic data analyses (Fig. S9B, (Wuest et al., 2010)). However, mCherry-AGO9 protein fusion expression in the egg cell does not agree with previously published results obtained by immuno-fluorescence (Olmedo-Monfil et al., 2010). Like mCherry-AGO8, the clade II-member mCherry-AGO2 is also specifically detected in the egg cell (Fig.1I) while clade II-member mCherry-AGO3 is solely detected in the chalazal seed coat of the ovule (Fig. 1J; (Jullien et al., 2020)). The last clade II-member, GFP-AGO7, is detected in all cell types of the female gametophyte and surrounding integument, with significant enrichment in the egg cell, similarly to AGO1 (clade I), AGO4 and AGO6 (clade III) (Fig. 1K).

Overall, our analysis shows that all ten *Arabidopsis* AGOs are detected in mature ovules before fertilization. Within the female gametophyte, their accumulation seems to be particularly enriched in the egg cell compared to the central cell. Preferential AGO expression in the egg cell was confirmed using previously published female gametophyte transcriptome data obtained by laser-capture microdissection (Fig. S9A,(Wuest et al., 2010)). Noteworthy, our data, together with the potential *ago8* phenotype previously described during MMC development (Hernández-Lagana et al., 2016), suggest that AGO8 might be functional. Nonetheless, *AGO8* is commonly thought to be a pseudogene, as its ORF displays a premature stop codon predicted to result in a truncated protein containing a PAZ domain but not the catalytic PIWI domain (Takeda et al., 2008). Our observation of mCherry-AGO8 protein specifically in the egg cell (Fig. 1G) could therefore reflect the accumulation of a truncated mCherry-AGO8 protein in this particular cell type. We could not test this hypothesis using western blotting, however, due to the very low expression levels of mCherry-AGO8. It is nevertheless tempting to hypothesize that a truncated AGO8 protein could act as a dominant-negative sRNA regulator specifically in the egg cell, as was shown for the dominant *ago5-4* allele which similarly encodes a truncated PAZ-proficient yet PIWI-deficient protein (Tucker et al., 2012). Taken together, our results uncover a complex sRNA-loading landscape likely granting functional diversity in the egg cell.

### AGO expression patterns in the male gametophyte

Analysis of the translational reporters in mature pollen grains (Fig. 2A-H) revealed a preferential enrichment of some AGOs in sperm cells. Indeed, mCherry-AGO1, mCherry-AGO2, GFP-AGO7, mCherry-AGO4 and mCherry-AGO9 were solely detected in those cells (Fig. 2 B, D,F-H). mCherry-AGO5 and mCherry-AGO6 were mainly detected in sperm cells, but also in the vegetative cell, albeit only at substantially lower levels (Fig. 2 C, E). AGO5 sperm cell expression was previously reported (Borges et al., 2011; Tucker et al., 2012). Signals from the mCherry-AGO3, mCherry-AGO8 and GFP-AGO10 reporters were not detected in mature pollen.

**Fig. 2.**
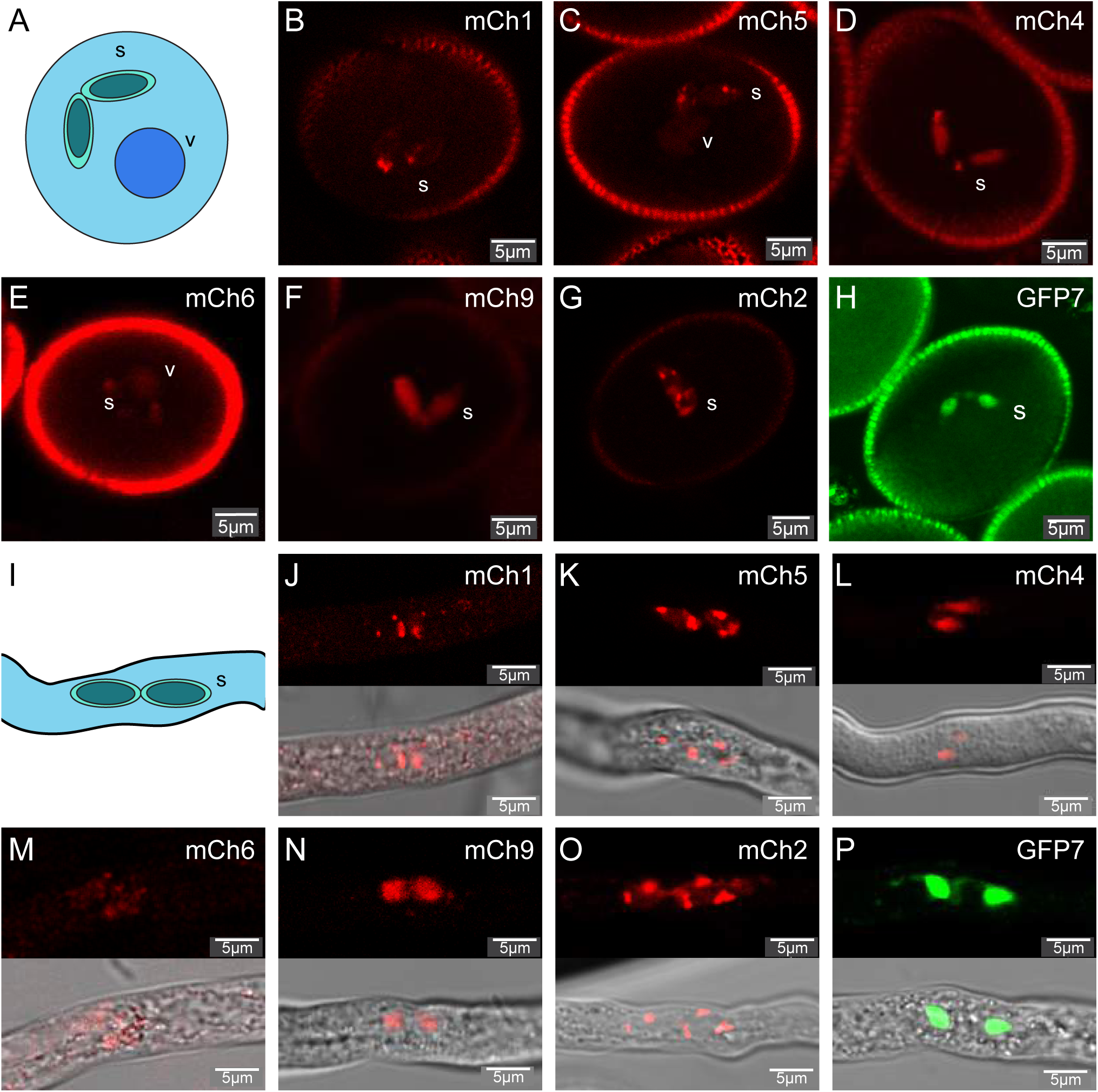
Argonaute expression in pollen. (A-H) AGOs in the mature male gametophyte. Schematic representation of a mature pollen grain of Arabidopsis thaliana illus-trating the two major cell type: the vegetative cell (v) and the two sperm cells (s). (B-H) Confocal images representing the expression of 7 Arabidopsis AGOs expressed in the pollen grain: mCherry-AGO1 (B), mCherry-AGO5 (C), mCherry-AGO4 (D), mCherry-AGO6 (E), mCherry-AGO9 (F), mCherry-AGO2 (G) and GFP-AGO7 (H). (I-P) Argonaute expression in germinating pollen tube. (I) Schematic representation of a growing pollen tube of Arabidopsis thaliana illustrating the two sperm cells (s). (J-P) Confocal images representing the 7 Arabidopsis AGOs expressed in in germinated pollen grain: mCherry-AGO1 (B), mCherry-AGO5 (C), mCherry-AGO4 (D), mCher-ry-AGO6 (E), mCherry-AGO9 (F), mCherry-AGO2 (G) and GFP-AGO7 (H). Scale bars represent 5 μm.

To address if paternally-expressed AGOs could be potentially transmitted during fertilization, we investigated the presence of the fusion proteins’ fluorescent signals in germinated pollen (Fig. 2I-P). All AGOs exhibiting sperm cells expression were indeed also detected in germinated pollen, suggesting that AGO-loaded sRNAs of paternal origin could be transported to the egg cell and potentially regulate gene expression in the zygote at, or shortly after, fertilization. Although such phenomena have been highlighted in the context of mammalian embryonic development (Conine et al., 2018; Sharma et al., 2018; Yuan et al., 2016), they have been seldom documented in plants, including in *Arabidopsis*. Indeed, most mutants affecting embryonic development are sporophytic recessive rather than showing a paternal gametophytic effect (Meinke, 2020), which is the case for mutant alleles of *DCL1*, encoding the main microRNA processing enzyme in *Arabidopsis* (Nodine and Bartel, 2010). Recently, however, a mutant allele of *MIR159* was reported to display paternal effects on endosperm development owing to the downregulation of the MYB33 transcription factor (Zhao et al., 2018b). The high expression of miR159 in sperm cell as well as the very early nature of the observed phenotype suggest indeed that miR159 could be delivered paternally at fertilization.

### AGO expression patterns in the early seed

To gain insights into AGO accumulation during early seed development, we observed fluorescent signals in seeds at one Day-After-Pollination (1 DAP), using confocal microscopy (Fig. 3A-I). At this stage, we could detect expression of eight out of ten *Arabidopsis* AGOs, while signals from mCherry-AGO2 and mCherry-AGO8 were below detection limit. Clade I mCherry-AGO1 is detected in both inner and outer integument layers in the sporophytic tissue as well as in the embryo, but is below detection in the endosperm (Fig. 3A). Similarly, clade I mCherry-AGO5 is detected in the embryo and inner (but not outer) integument, while it is excluded from the endosperm (Fig. 3B). Despite not being detected in the egg cell, clade I GFP-AGO10 could be detected in the early embryo but not in the endosperm (Fig. 3C). Strong GFP-AGO10 accumulation is also observed in the funiculus’ vasculature and the seed/ovule vascular termination (Fig. S7). As previously described, accumulation of the clade II mCherry-AGO3 is limited to the chalazal seed coat (Fig. 3H; (Jullien et al., 2020)). Clade II GFP-AGO7 is detected in all cell types except the endosperm, and its accumulation is particularly strong in the chalazal seed coat (Fig. 3I).

**Fig. 3.**
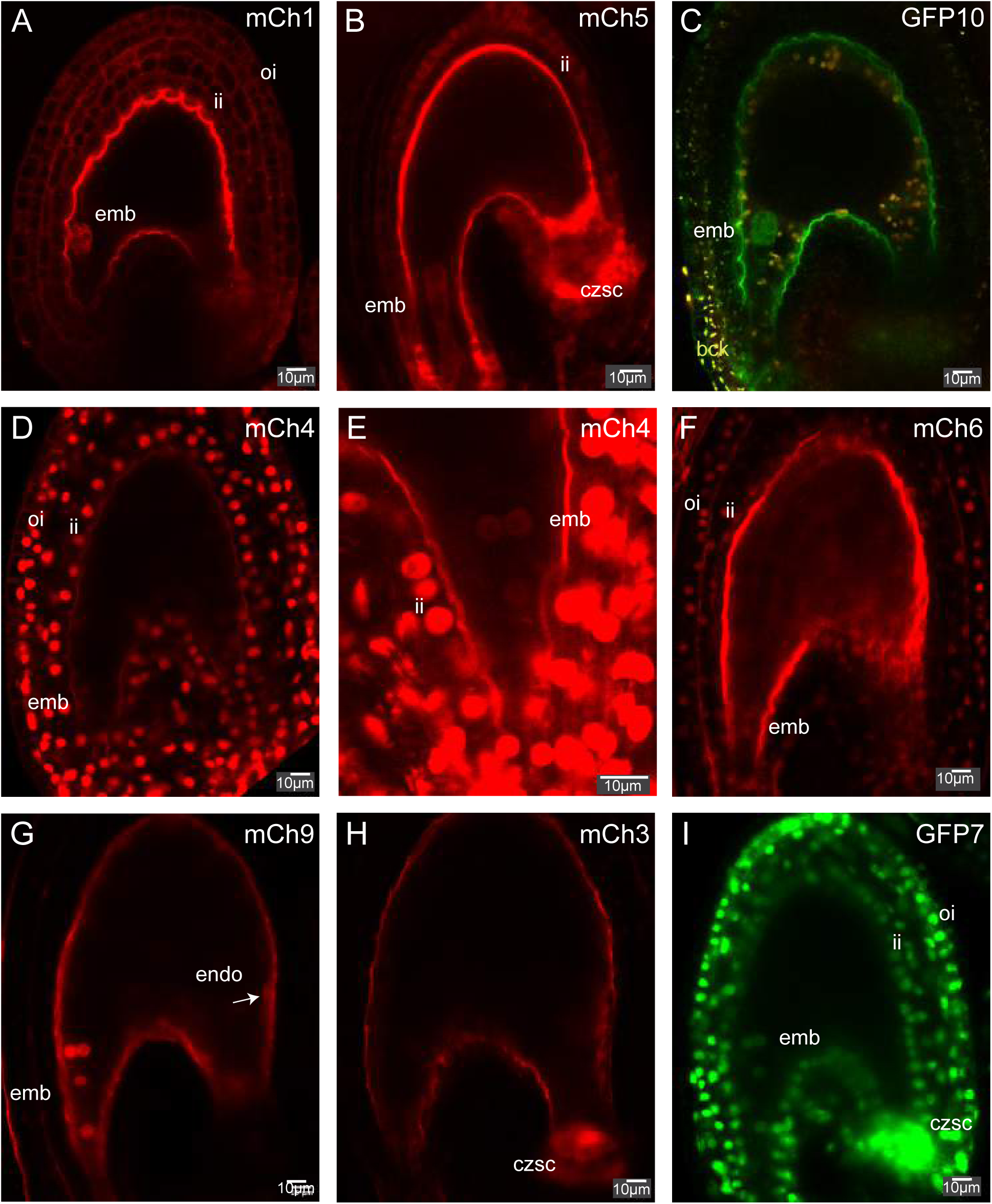
Argonaute expression in developing seed. (A-I) Confocal images representing Arabidopsis AGOs expressed in the developing seeds at 2 day after pollination: mCherry-AGO1 (A), mCherry-AGO5 (B), GFP-AGO10 (C), mCherry-AGO4 (D-E), mCherry-AGO6 (F), mCherry-AGO9 (G), mCherry-AGO3 (H) and GFP-AGO7 (I). Scale bars represent 10 μm. Abbreviation: emb (embryo), endo (endosperm), inner integument (ii), outer integuments (oi), Chalazal seed coat (czsc).

Among the clade II AGOs involved in RdDM, mCherry-AGO4 shows the strongest accumulation, especially in the sporophytic tissues (Fig. 3D). mCherry-AGO4 can be detected in the early embryo (Fig 3E) but below detection in the endosperm. The expression pattern of mCherry-AGO6 is similar to that of mCherry-AGO4 (Fig.3F), although the signal is of lower intensity. This result agrees with the known redundancy of AGO4 and AGO6 in mediating DNA methylation and TGS at some genetic loci (Stroud et al., 2012; Zheng et al., 2007). mCherry-AGO9 could be detected in the embryo and also in the endosperm, although not in the integuments (Fig. 3G). In fact, mCherry-AGO9 is visible from the first nuclear division of the endosperm, upon which its signal decreases in intensity, although it is still detected at the 4-cells embryo stage (Fig. 3G and Fig. S8B-C). Based on our reporter constructs, AGO9 appears therefore, to be the only AGO detectable during endosperm development.

To conclude, we observe a strong asymmetry of AGOs’ patterns between the endosperm and the embryonic lineages. This difference is supported by LCM transcriptomic data from developing seeds (Fig. S8, (Belmonte et al., 2013)) and suggests a less active involvement of RNA silencing pathways in the endosperm compared to the embryo, during early seed development. In *Arabidopsis*, the RdDM pathway maintains cytosine DNA methylation in the CHH context (where H is any nucleoside but G), which cannot be perpetuated by maintenance DNA methylases. This process relies on the constant action of *de novo* DNA methyltransferases (DRM1 and DRM2) that are guided to their cognate target loci by interacting with siRNA-loaded, Class II AGOs. Similarly to what was reported for the *de novo* DNA methyltransferases (Jullien et al., 2012), the asymmetric expression of Class II AGOs observed between the embryonic and endosperm lineages could contribute to the high CHH methylation observed in the former compared to the latter (Bouyer et al., 2017; Gehring et al., 2009; Hsieh et al., 2009; Kawakatsu et al., 2017).

### AGO expression patterns in heart-stage embryo

In order to investigate AGO accumulation patterns in the differentiated zygote, we dissected heart-stage embryos in which most early developmental decisions have already taken place, and performed confocal imaging using the fluorescently tagged AGO transgenic lines (Fig. 4A-H). As previously reported (Du et al., 2019; Lynn et al., 1999), clade I mCherry-AGO1 is expressed in all cells of the heart-stage embryo (Fig. 4B). Similarly, we observed a ubiquitous expression of the clade II mCherry-AGO4 and mCherry-AGO6 (Fig. 4E-F). By contrast, clade III mCherry-AGO2, mCherry-AGO3 and clade II mCherry-AGO8 could not be detected in the heart-stage embryo. Clade I GFP-AGO10 displayed its previously reported pattern (Du et al., 2019; Tucker et al., 2008), with fluorescent signals observed in the adaxial part of cotyledons and in the pre-vasculature (Fig 4D).

**Fig. 4.**
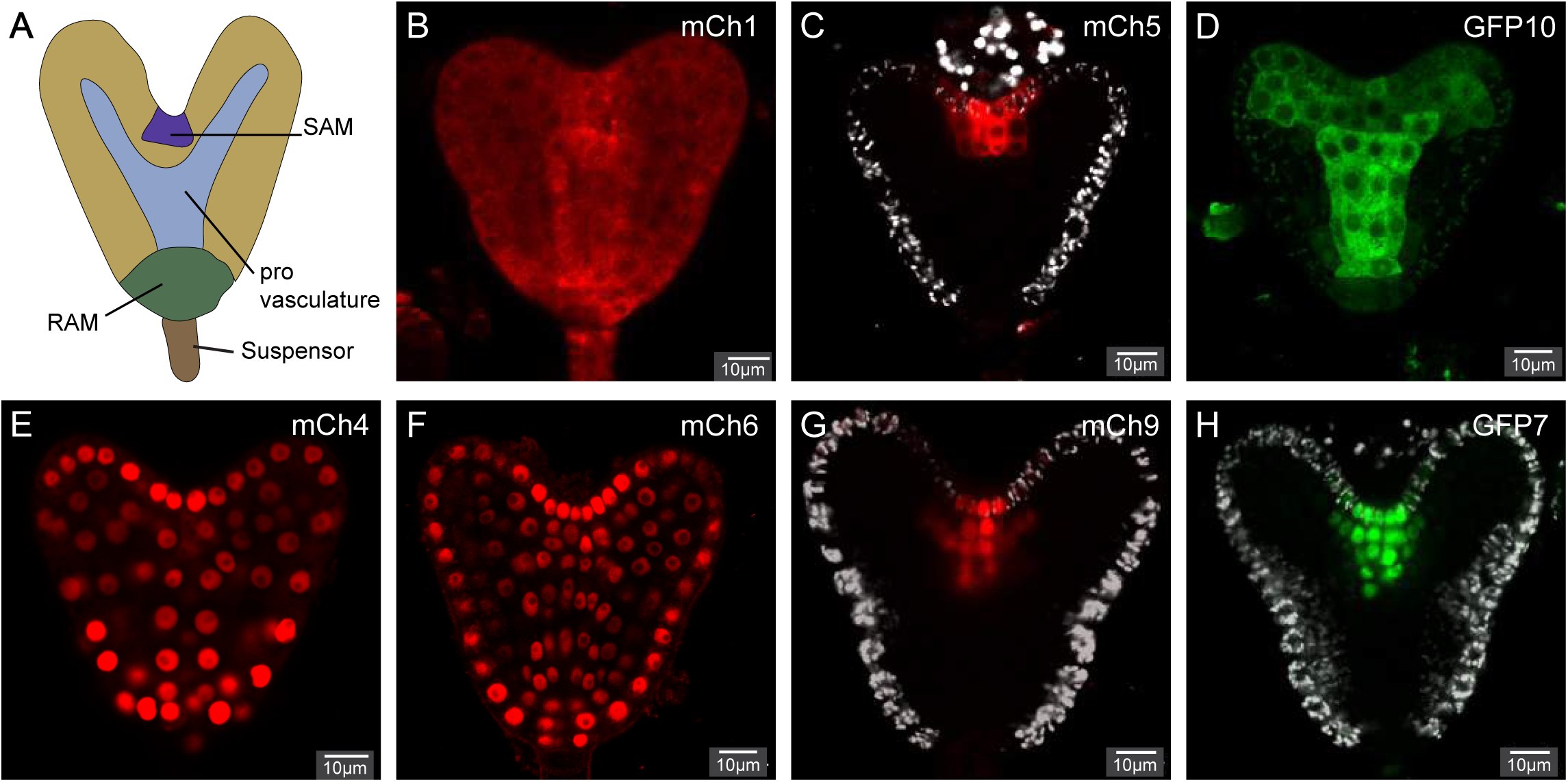
Argonaute expression in heart stage embryo. (A) A simplified representation of a heart stage embryo illustrating the different cell types. (B-H) Confocal images representing the 7 Arabidopsis AGOs expressed in heart stage embryo: mCherry-AGO1 (B), mCherry-AGO5 (C), GFP-AGO10 (D), mCherry-AGO4 (E), mCherry-AGO6 (F), mCherry-AGO9 (G), and GFP-AGO7 (H). Scale bars represent 10 μm.

The three remaining AGOs, AGO5 (clade I), AGO9 (clade III) and AGO7 (clade II) accumulate specifically in the shoot apical meristem region (SAM) of the heart-stage embryo (Fig. 4C, G, H), a pattern already documented for AGO5 using a YFP fluorescent reporter (Tucker et al., 2012). In agreement with our results, AGO5, AGO7, and AGO9 transcript were both found to be enriched in meristematic stem cells at the embryonic stage (Gutzat et al., 2018). Although AGO7 and AGO9 functions in meristems remain to be investigated, recent work suggests that AGO5 is involved in regulating flowering, perhaps via a floral meristem-specific activity (Roussin-Léveillée et al., 2020). Indeed, *ago5* mutants display an early flowering phenotype without overtly altered leaf morphology (Fig.S4, (Roussin-Léveillée et al., 2020)). This effect on flowering is thought to proceed through AGO5-miR156 specific interactions repressing, in turn, accumulation of SPL transcription factors. However, miR156 was previously shown to affect both flowering and leaf morphology (Wu et al., 2009), thereby suggesting a more complex underpinning to the *ago5* phenotype. This possibly involves differential AGO loading and functions for miR156 or highly sequence-related miR157 isoforms in meristematic regions (Ebhardt et al., 2010; He et al., 2018).

### AGO intracellular localization patterns

*Arabidopsis* AGO1 and AGO4 have been shown to shuttle between the cytoplasm and the nucleus although their respective steady-state subcellular localizations seem to reflect their involvement in either PTGS (clade I and II) and TGS/RdDM (Clade III),: AGO1 is mostly cytoplasmic and AGO4 nuclear (Bologna et al., 2018; Ye et al., 2012). In agreement with their involvement in Clade I and II, mCherry-AGO1, mCherry-AGO5, mCherry-AGO2, mCherry-AGO3, GFP-AGO10 are mostly localized in the cytoplasm in reproductive cells (Fig. 1B-D, Fig.1I-J, Fig.3A-C, Fig3H, Fig. 4B-D). Their cytoplasmic localization is consistent with studies showing their association with the translation machinery (Jullien et al., 2020; Marchais et al., 2019). However, we observed that mCherry-AGO1, mCherry-AGO5, mCherry-AGO2 form cytosolic aggregates observed only certain cell types such as the nucellus and sperm cells (Fig. 1B-C and Fig. 2B-C, G). Nonetheless, caution should be exerted in interpreting these observations given that artefactual aggregation has been reported with mCherry-tagged proteins (Cranfill et al., 2016; Landgraf et al., 2012). Since it is likely that these AGO foci occur in tissues where AGO1, AGO5 and AGO2 are particularly highly expressed, further work will be required to assess if mCherry-AGO1, mCherry-AGO5, and mCherry-AGO2 aggerates represent relevant biological entities.

The main Clade II AGOs, mCherry-AGO4 and mCherry-AGO6, are localized in the nucleus in all reproductive cell types analyzed (Fig. 1E-F, Fig. 2D-E, Fig. 3D-F, Fig. 4E-F). Occasionally, cytoplasmic localization could be observed in the ovule integument or embryo likely due to the disruption of the nuclear membrane during cell division. Nuclear localization for AGO4 and AGO6 was previously observed in other tissues (Ye et al., 2012; Zheng et al., 2007). Perhaps more strikingly, mCherry-AGO9 displays nuclear localization in somatic tissues (Fig. 1H, Fig. 3G, Fig. 4G, Fig. S8) but appears to be also partially localized to the cytoplasm in the central and egg cells (Fig. 1H). Combined cytoplasmic and nuclear localizations of AGO9 were previously observed by immunolocalization in ovule primordia (Rodríguez-Leal et al., 2015; Zhao et al., 2018a). However, in those studies, AGO9 was found to accumulate in cytoplasmic foci which we could not observe with our reporter construct in the tissues examined. The mCh-AGO8 translational reporter, unlike other RdDM AGOs, was mainly localized to the cytoplasm of the egg cell (Fig. 1G). As discussed above, this unusual intra-cellular localization for a Clade II AGO could result from AGO8 being a truncated, perhaps dominant-negative, protein.

One of the most intriguing intracellular localization patterns was that of GFP-AGO, found mainly localized to the nucleus. However, in some cells of the integument, in the egg cell and in the central cell, clear cytosolic localization was additionally observed, with the presence of cytoplasmic foci (Fig. 1K-L). Localization of GFP-AGO7 to cytoplasmic foci named “sRNA bodies” was previously observed in *Nicotiana benthamiana* leaves during transient overexpression of GFP-AGO7 (Jouannet et al., 2012). Nuclear GFP-AGO7 accumulation was not reported in this study. Several reasons could underpin this discrepancy, such as the difference in the promoter used (*p35S* versus cognate *pAGO7* promoters) and also, importantly, the tissues analyzed (mature leaves or roots *versus* reproductive tissue). Interestingly, retention of AGO7 in the nucleus, using an NLS signal-peptide fused to AGO7 in stable *Arabidopsis* transformant, did not complement the so-called leaf zippy phenotype of *ago7* mutant (Hunter et al., 2003; Jouannet et al., 2012). This result shows that AGO7 cytoplasmic localization is necessary for its function during leaf development. Similarly to AGO7, it was shown that nuclear retention of AGO1 does not allow complementation of the *ago1* mutant phenotype as AGO1 shuttling between the nucleus and the cytoplasm is required for its function in the *Arabidopsis* miRNA pathway (Bologna et al., 2018). The AGO7 zippy mutant phenotype relies on the compromised loading, by AGO7, and function of miR390. The fact that our construct rescues the *ago7* zippy leaf phenotype (Fig. S6) and displays both nuclear and cytoplasmic localization in the inspected tissues suggests that AGO7, similarly to AGO1, shuttles between both subcellular compartments in a manner possibly developmentally regulated. Nonetheless, nuclear AGO7 might have additional, as yet uncharacterized functions in reproductive tissues, as shown recently for nuclear AGO1 (Liu et al., 2018).

To conclude, we have generated full locus N-terminal tagged fluorescent constructs of all *Arabidopsis* AGOs under their native endogenous promoter. Though we have analyzed their expression pattern and intracellular localization in a study focused primarily on reproductive tissues, there is every reason to believe that these constructs and transgenic lines will be equally useful to study AGOs regulations in other developmental contexts or during stress. Fluorescent AGO-reporters have been sporadically described in *Arabidopsis*, yet they are often marred by biological incongruities such as the use of overexpression promoters or C-terminal fusions known to affect AGO functions. The uniform set of tools presented in this study might help bridging this still significant gap across the plant RNA silencing community.

## Materials and methods

### Plant Material and Growth Conditions

After three days at 4°C in the dark, seeds were germinated and grown on soil. Plants were grown under long days at 20-21°C (16h light/8h night). All plants were in Columbia (Col-0) accession. The mutants and lines described in this work correspond to the following: *ago1-36* (SALK_087076,(Baumberger and Baulcombe, 2005)), *ago2-1* (salk_003380,(Lobbes et al., 2006)), *ago3-3* (GABI-743B03,(Jullien et al., 2020)), *ago4-5* (WiscDsLox338A0, (Stroud et al., 2012)), *ago5-1* (salk_063806, (Katiyar-Agarwal et al., 2007)), *ago6-2* (salk_031553, (Zheng et al., 2007)), *ago7-1* (salk_037458, (Vazquez et al., 2004)), *ago8-1* (salk_139894, (Takeda et al., 2008), *ago9-1* (salk_127358, (Katiyar-Agarwal et al., 2007)) and *ago10-1* (SALK_000457). The insertion lines were provided by The Nottingham *Arabidopsis* Stock Centre (NASC) (http://arabidopsis.info/). Pollen were germinated in Pollen growth medium at 21°C in the dark 5 hours to over-night (Hamamura et al., 2011).

### Microscopy

Fluorescence images were acquired using laser scanning confocal microscopy (Zeiss LSM780 or Leica SP5). Brightness was adjusted using ImageJ (http://rsbweb.nih.gov/ij/) and assembled using ImageJ or Adobe Illustrator.

### Plasmid Construction and Transformation

All DNA fragments were amplified by PCR using the Phusion High-Fidelity DNA Polymerase (Thermo). Primer sequences can be found in Supplementary Table S1. All constructs were generated using Multisite Gateway technology (Invitrogen). *A. thaliana* transformation was carried out by the floral dip method (Clough and Bent, 1998). All plasmids were transformed into Col-0 and their respective mutants and for mCherry-AGO6 also in LIG1-GFP marker line (Andreuzza et al., 2010). Six to nineteen transgenic lines (T1) were analysed and showed a consistent fluorescence expression pattern using a Leica epifluorescence microscope or a Leica SP5. One to three independent lines with single insertions, determined by segregation upon BASTA selection, were used for further detailed confocal analysis.

## Author contributions

PEJ conceived the study. PEJ, NP and JAS generated the transgenic lines. PEJ and DMVB performed the imaging. DMVB and JAS tested the complementation. PEJ wrote the manuscript, which was further edited and amended by OV.

## Acknowledgements

We would like to thank the following people for their help: Andre Imboden and Jasmine Sekulovski for support in plant growth, Nicolas G. Bologna for critical reading of the manuscript We would like to thank the ETH Scope M and Microscopy Imaging Center of the University of Bern.

## Data availability

Plasmids and lines used in this study will be made available to the community.

## Supplementary information

Additional Supporting Information may be found in the online version of this article

---

**Fig. S1.**
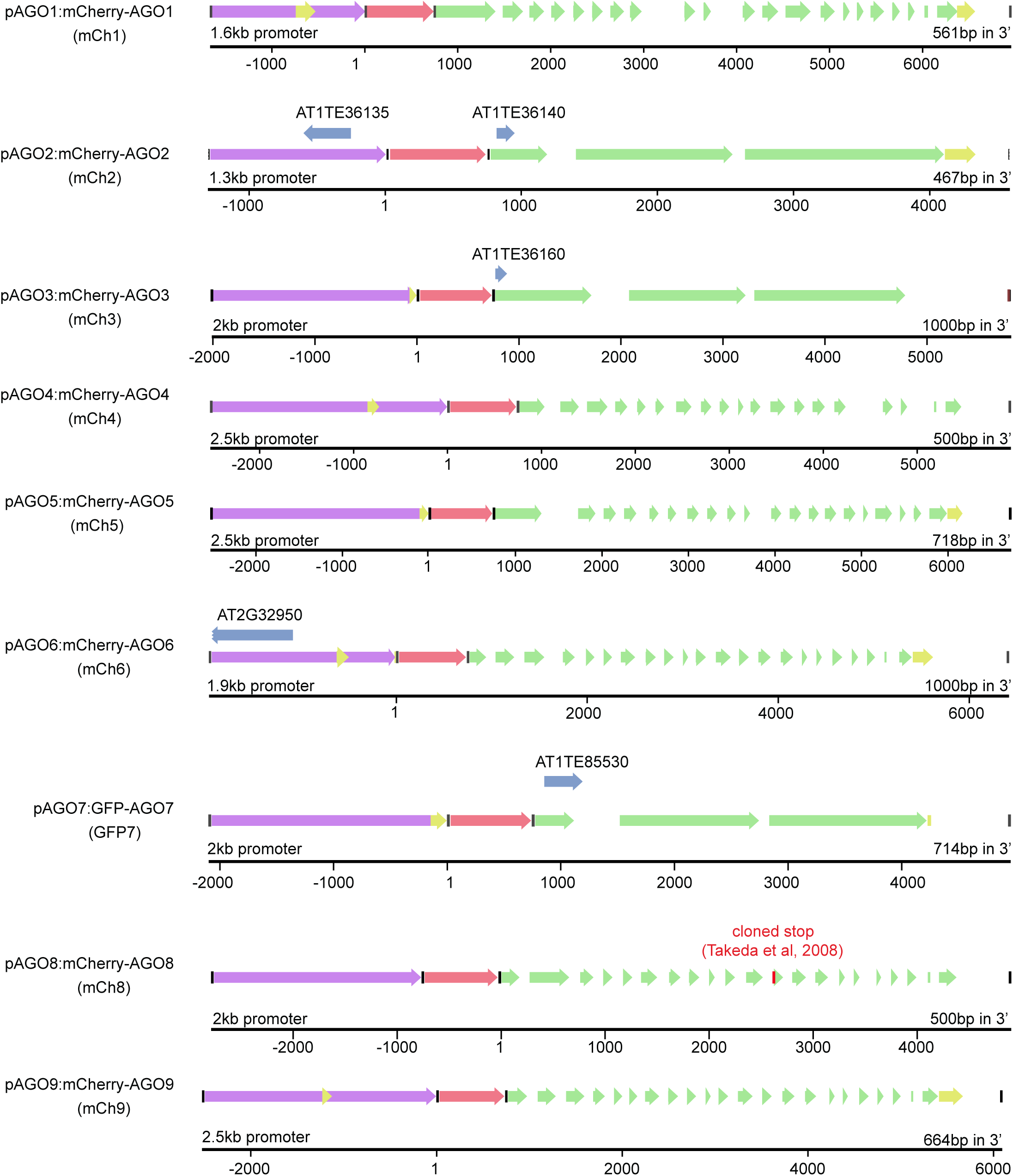
Schematic representation of the construct used in this study. The different features are represented by arrows: promoter (purple), UTR (yellow), fluorescent protein (red), exon (green) and additional annotation (blue).

**Fig. S2.**
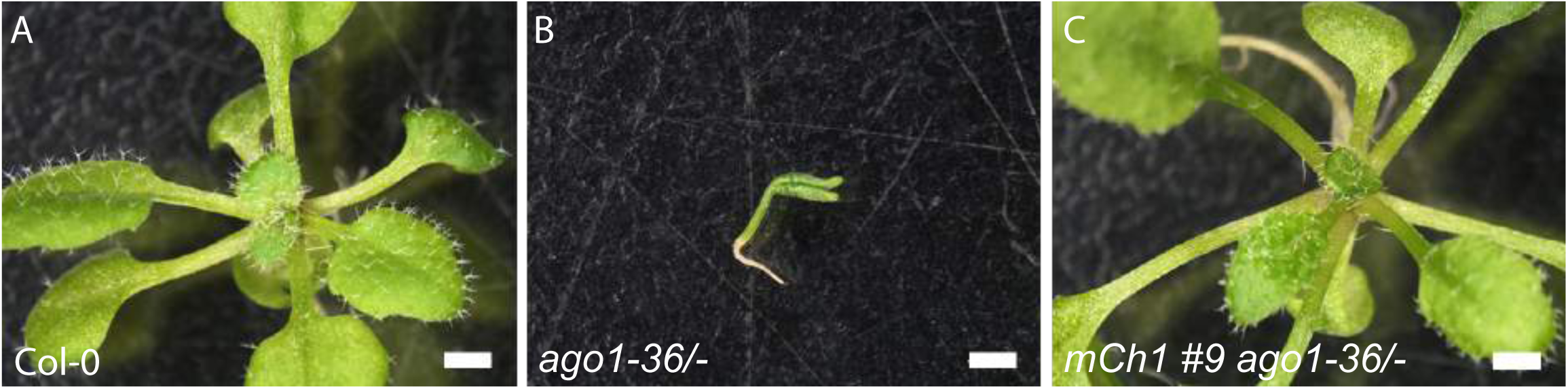
Complementation of ago1-36 mutant by pAGO1:mCherry-AGO1. Illustrative pictures showing the lack of developmental phenotype in Col-0 (A) and mCh1#9 ago1-36/-(C) compared to ago1-36/-. scale bar represents 1mm.

**Fig. S3.**
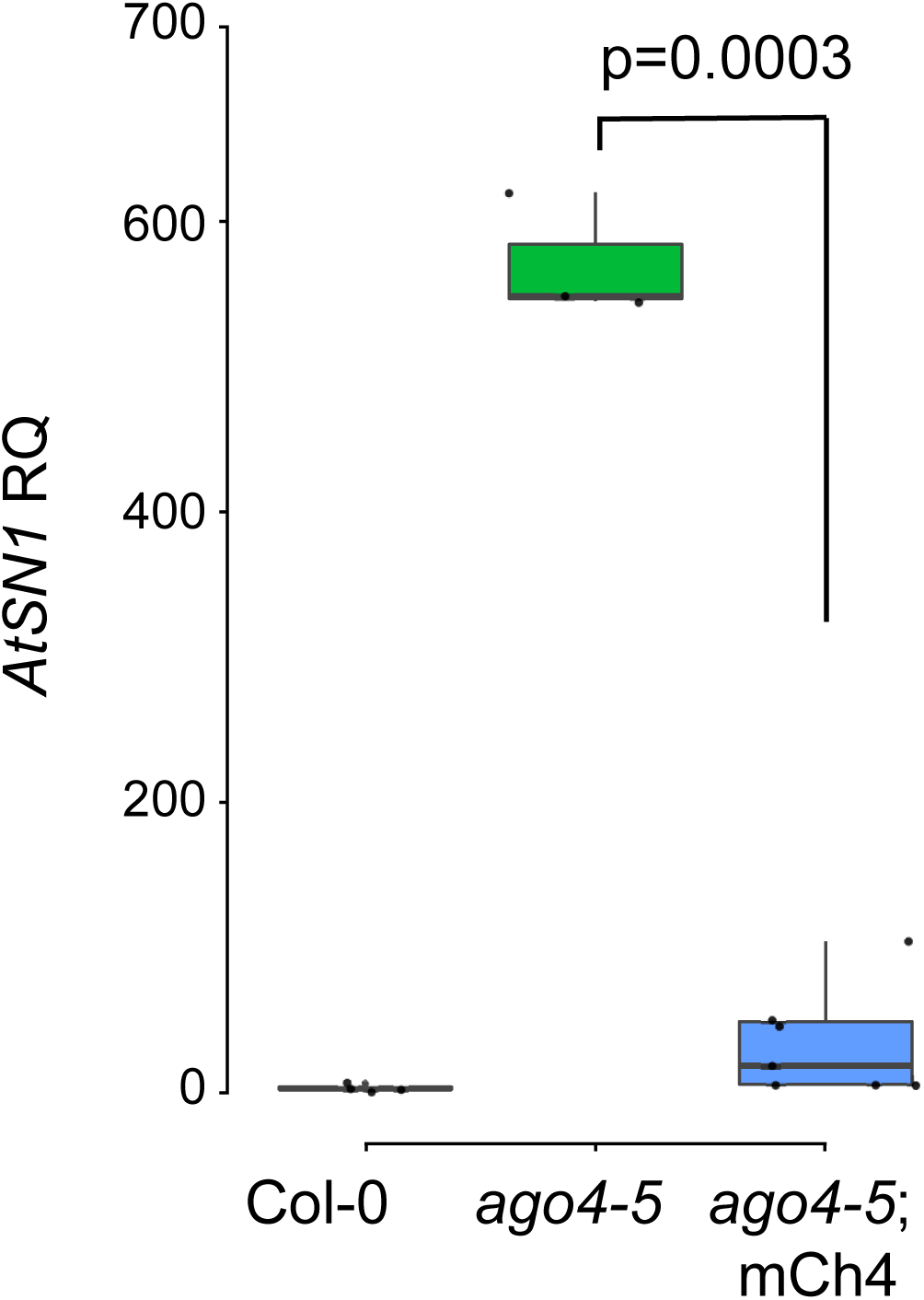
Complementation of ago4-5 mutant by pAGO4:mCherry-AGO4. qPCR showing the absence of AtSN1 ectopic expression in rosette leaves of seven independent lines expressing the construct pAGO4:mCherry-AGO4 (mCh4) in ago4-5 background compared to Col-0 and ago4-5. Actin2 was used as endogenous control. p indicates the p value obtained by a Student’s T-Test.

**Fig. S4.**
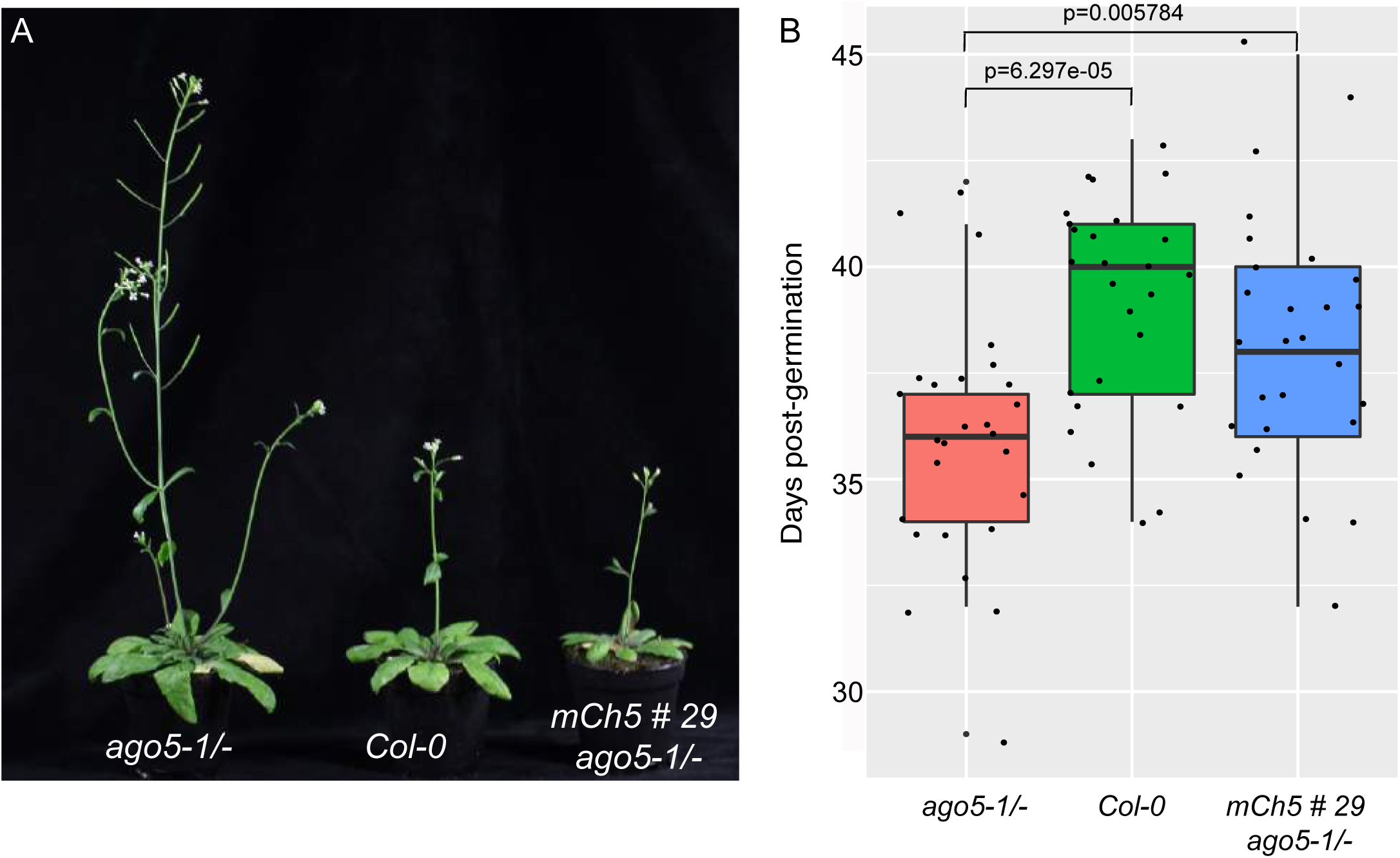
Complementation of ago5-1 mutant by pAGO5:mCherry-AGO5. (A) Illustrative pictures of the early flowering phenotype of ago5-1/- compared to Col-0 and mCh5 #29 ago5-1/-. (B) Quantification of ago5-1/- flowering phenotype in ago5-1/- (n=27), Col-0 (n=26) and mCh5#29 ago5-1/- (n=27). p indicates the p value obtained by a Student’s T-Test.

**Fig. S5.**
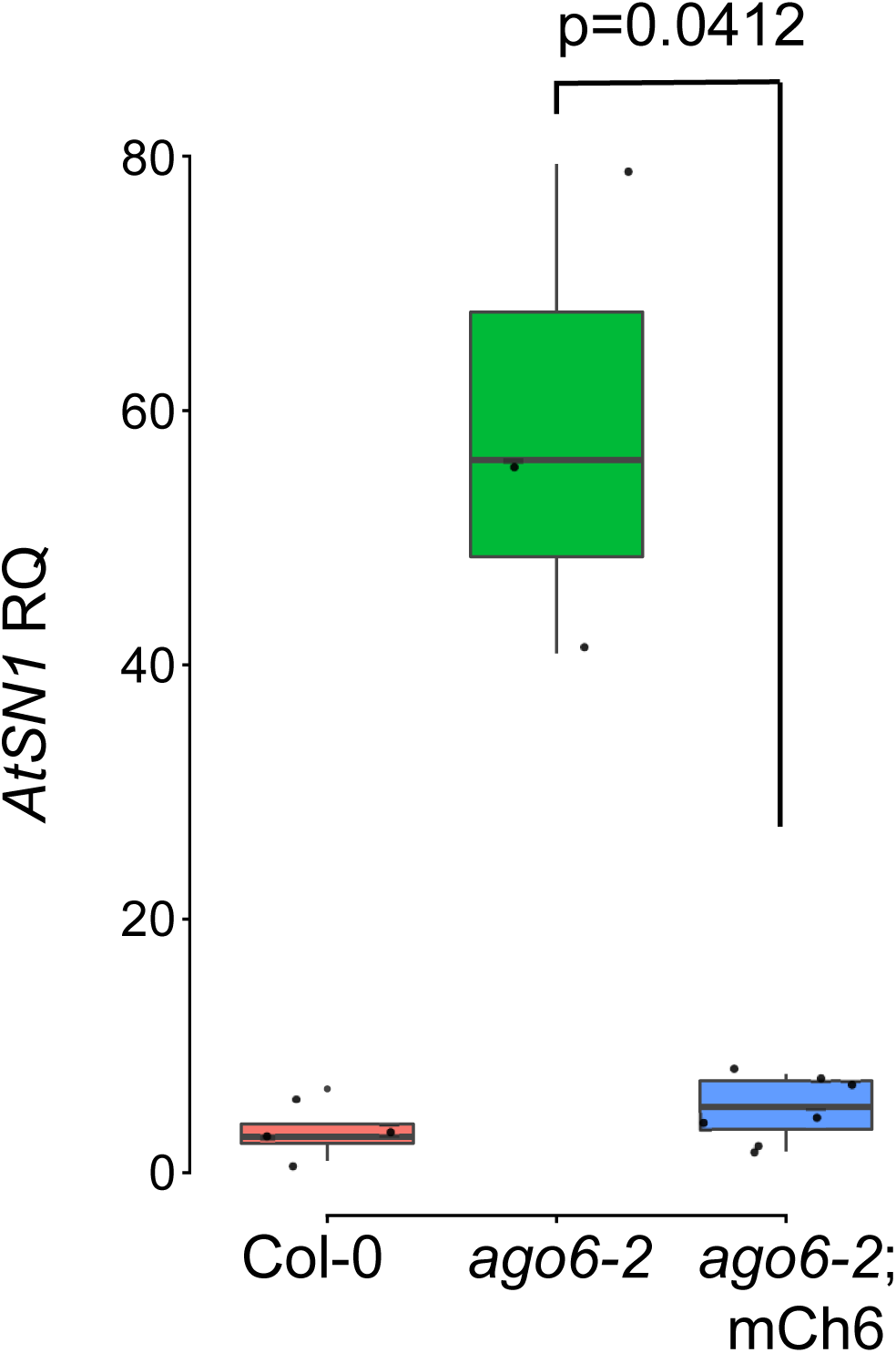
Complementation of ago6-2 mutant by pAGO6:mCherry-AGO6. qPCR showing the absence of AtSN1 ectopic expression in rosette leaves of seven independent lines expressing the construct pAGO6:mCherry-AGO6 (mCh4) in ago6-2 background compared to Col-0 and ago6-2. Actin2 was used as endogenous control. p indicates the p value obtained by a Student’s T-Test.

**Fig. S6.**
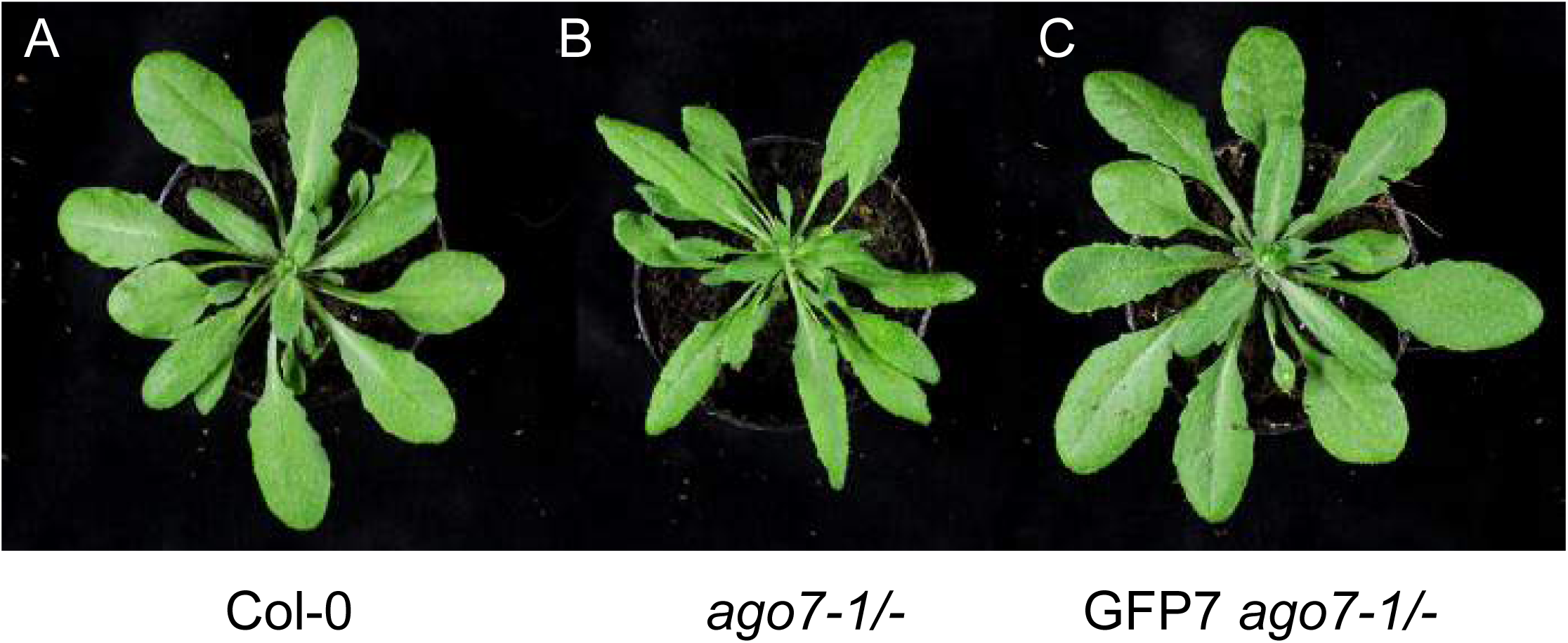
Complementation of ago7-1 mutant by pAGO7:GFP-AGO7. Illustrative pictures of the leaf phenotype of ago7-1/- (B) compared to Col-0 (A) and GFP-AGO7 ago7-1/- (C).

**Fig. S7.**
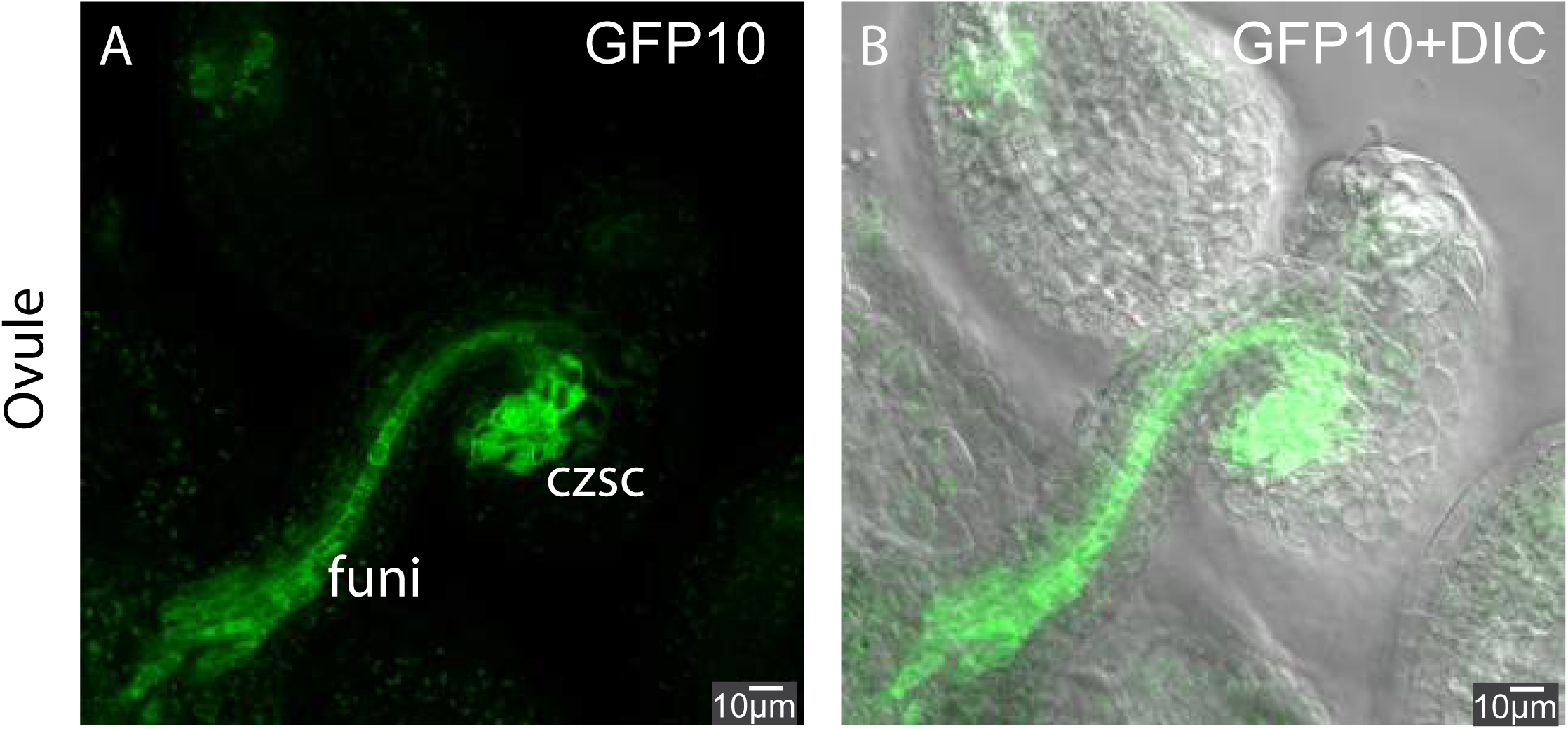
Additional pictures of pAGO10:GFP-AGO10. (A-B) Picture representing the expression in ovules of GFP-AGO10 in the vascular tissue of the funiculus (funi) and at the vascular termination in the chalazal seed coat (czsc). Scale bars represent 10μm.

**Fig. S8.**
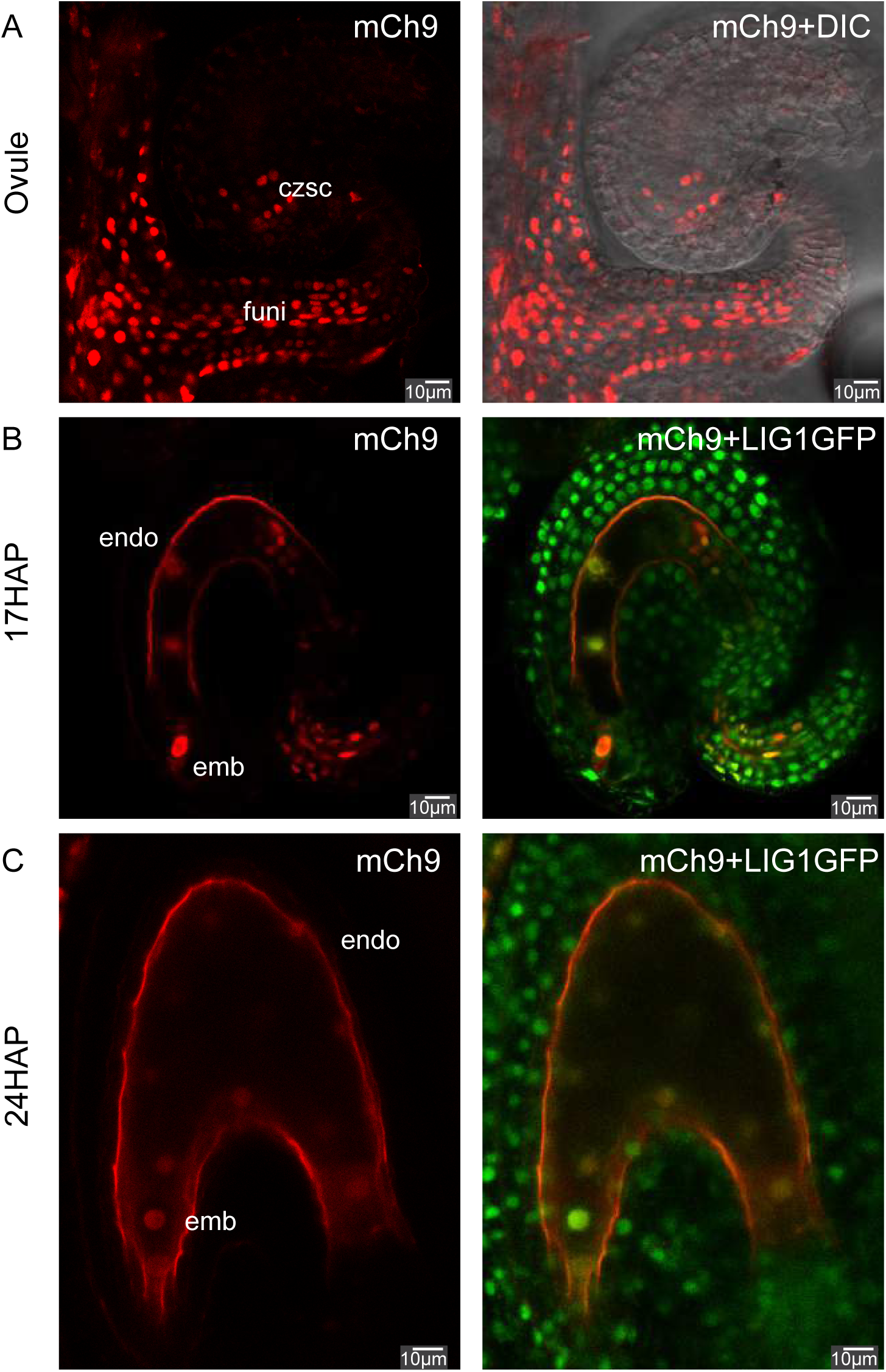
Additional pictures of pAGO9:mCh-AGO9. (A) Pictures representing the expression in ovules of mCherry-AGO9 in the funiculus (funi) and in the chalazal seed coat (czsc). (B-C) Pictures representing the expression in developing seeds of mCh9 in early embryo and endosperm at 17 hours after pollination (HAP) (B) and at 24 HAP (C). Scale bars represent 10 μm.

**Fig. S9.**
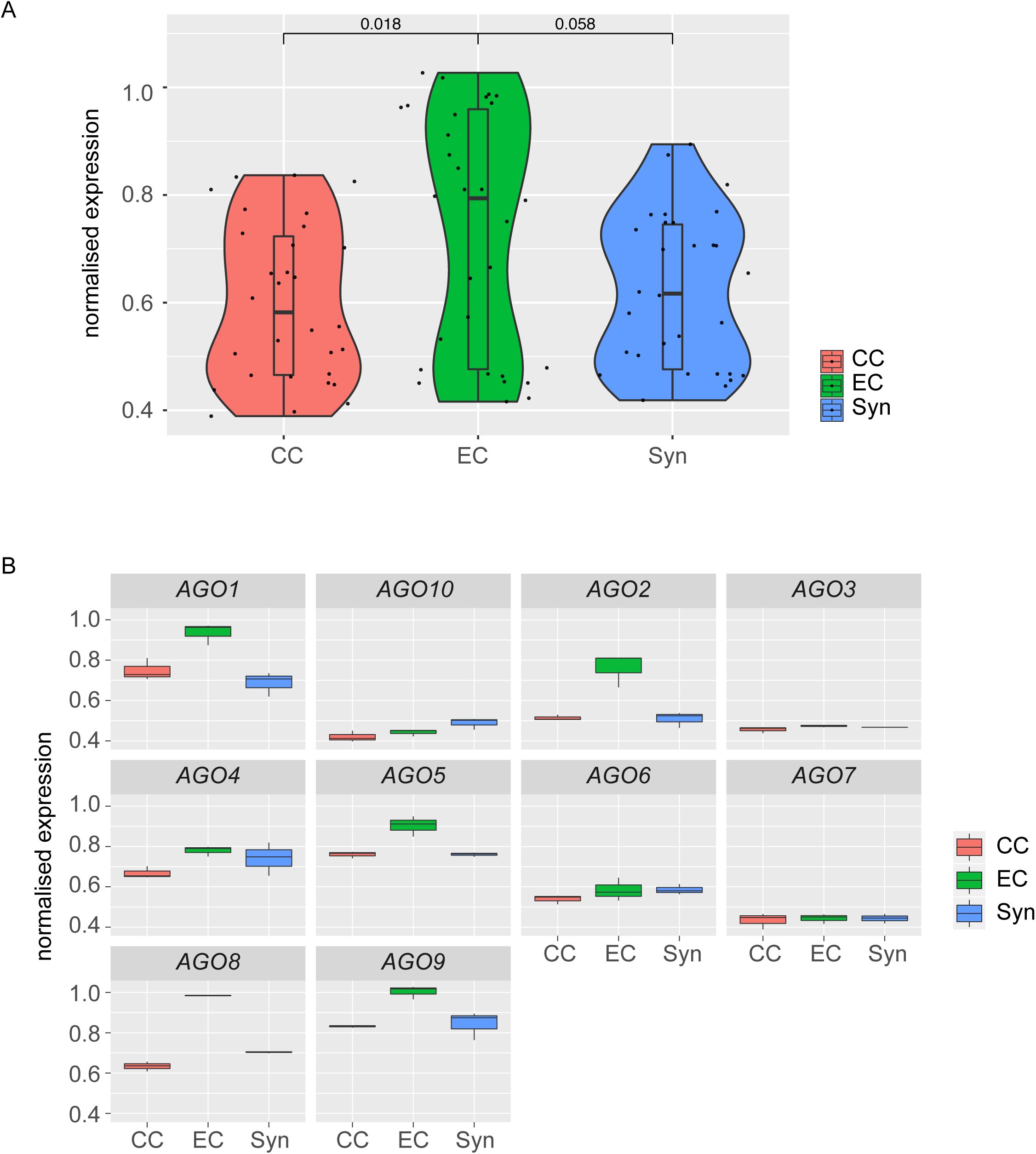
Arabidopsis Argonautes expression extracted from microarray data of LCM dissected female game-tophyte (Wuest et al, 2010) confirming the general enrichment of AGO transcript in the egg cell (EC) compared to central cell (CC) or synergids (Syn). (A) Violin plot representing the general enrichment of AGO expression in the EC. (B) AGOs individual expression boxplot in the different cell type. p values of a Wilcoxon test are indicated.

**Fig. S10.**
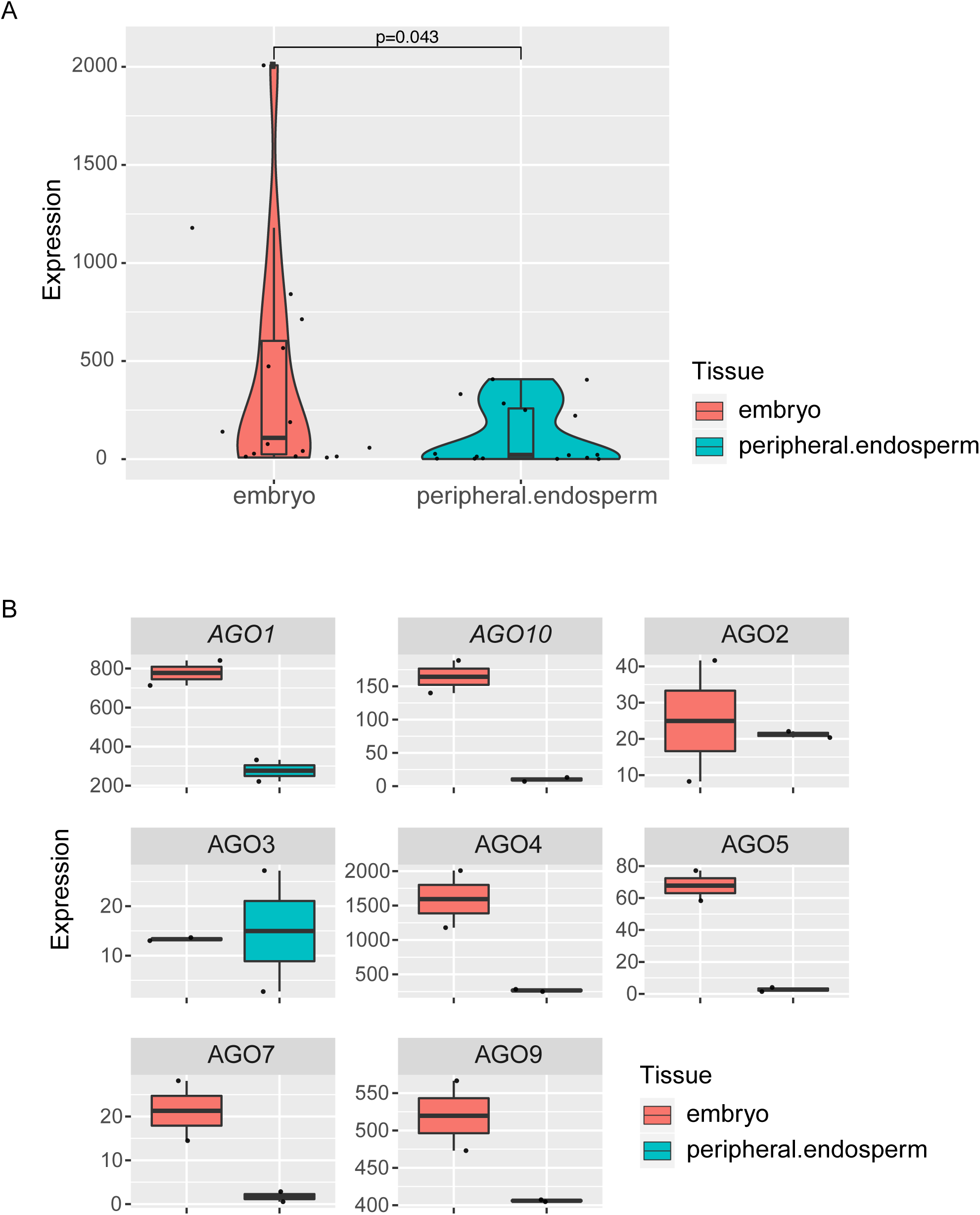
Arabidopsis Argonautes expression extracted from microarray data of LCM dissected seeds at the pre-globular stage (Belmonte et al., 2013) confirming the general enrichment of AGO transcripts in the embryo compared to the peripheral endosperm. (A) Violin plot representing the general enrichment of AGOs expression in the embryo. (B) AGOs individual expression boxplot in the different cell type. p values of a Wilcoxon test are indicated. AGO6 and AGO8 probes are not present in these data.

**Fig S11.**
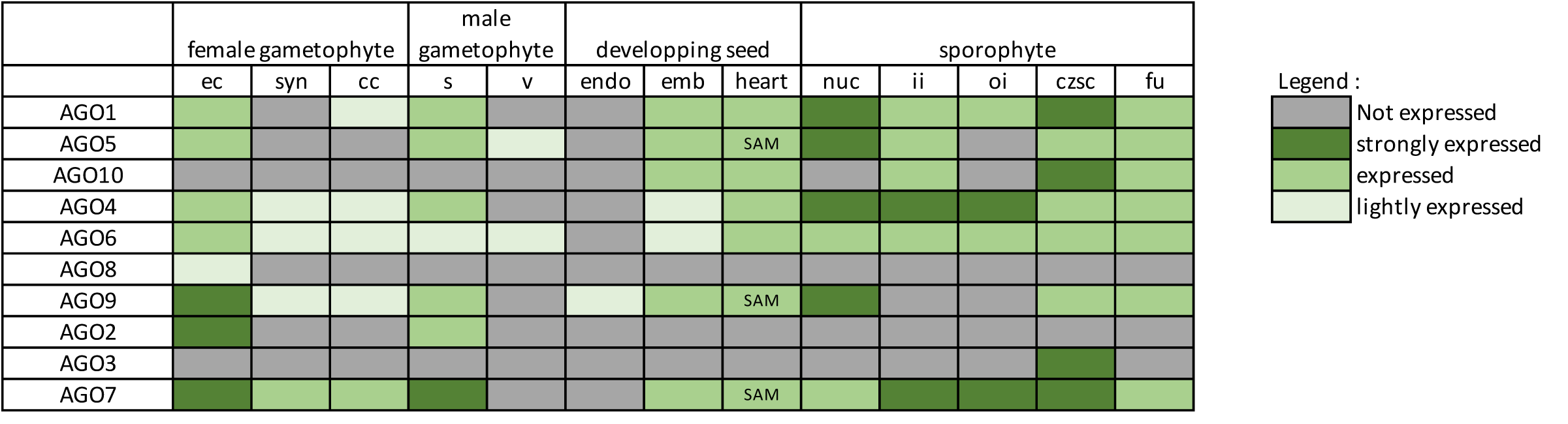
Summary table of AGO’s expression pattern in reproductive tissues.

## REFERENCES

Andreuzza, S., Li, J., Guitton, A.-E., Faure, J.-E., Casanova, S., Park, J.-S., Choi, Y., Chen, Z. and Berger, F. (2010). DNA LIGASE I exerts a maternal effect on seed development in Arabidopsis thaliana. Development 137, 73–81.

Baumberger, N. and Baulcombe, D. C. (2005). Arabidopsis ARGONAUTE1 is an RNA Slicer that selectively recruits microRNAs and short interfering RNAs. Proc. Natl. Acad. Sci. U. S. A. 102, 11928–11933.

Belmonte, M. F., Kirkbride, R. C., Stone, S. L., Pelletier, J. M., Bui, A. Q., Yeung, E. C., Hashimoto, M., Fei, J., Harada, C. M., Munoz, M. D., et al. (2013). Comprehensive developmental profiles of gene activity in regions and subregions of the Arabidopsis seed. Proc. Natl. Acad. Sci. 110, E435–E444.

Bologna, N. G. and Voinnet, O. (2014). The diversity, biogenesis, and activities of endogenous silencing small RNAs in Arabidopsis. Annu. Rev. Plant Biol. 65, 473–503.

Bologna, N. G., Iselin, R., Abriata, L. A., Sarazin, A., Pumplin, N., Jay, F., Grentzinger, T., Dal Peraro, M. and Voinnet, O. (2018). Nucleo-cytosolic Shuttling of ARGONAUTE1 Prompts a Revised Model of the Plant MicroRNA Pathway. Mol. Cell 69, 709-719.e5.

Borges, F. and Martienssen, R. A. (2015). The expanding world of small RNAs in plants. Nat. Rev. Mol. Cell Biol. 16, 727–741.

Borges, F., Pereira, P. a, Slotkin, R. K., Martienssen, R. a and Becker, J. D. (2011). MicroRNA activity in the Arabidopsis male germline. J. Exp. Bot. 62, 1611–20.

Borges, F., Parent, J. S., Van Ex, F., Wolff, P., Martínez, G., Köhler, C. and Martienssen, R. A. (2018). Transposon-derived small RNAs triggered by miR845 mediate genome dosage response in Arabidopsis. Nat. Genet. 50, 186–192.

Bouyer, D., Kramdi, A., Kassam, M., Heese, M., Schnittger, A., Roudier, F. and Colot, V. (2017). DNA methylation dynamics during early plant life. Genome Biol. 18, 179.

Carbonell, A., Fahlgren, N., Garcia-Ruiz, H., Gilbert, K. B., Montgomery, T. a, Nguyen, T., Cuperus, J. T. and Carrington, J. C. (2012). Functional analysis of three Arabidopsis ARGONAUTES using slicer-defective mutants. Plant Cell 24, 3613–29.

Castel, S. E. and Martienssen, R. A. (2013). RNA interference in the nucleus: Roles for small RNAs in transcription, epigenetics and beyond. Nat. Rev. Genet. 14, 100–112.

Clough, S. J. and Bent, A. F. (1998). Floral dip: A simplified method for Agrobacterium-mediated transformation of *Arabidopsis thaliana*. Plant J. 16, 735–743.

Conine, C. C., Sun, F., Song, L., Rivera-Pérez, J. A. and Rando, O. J. (2018). Small RNAs Gained during Epididymal Transit of Sperm Are Essential for Embryonic Development in Mice. Dev. Cell 46, 470-480.e3.

Cranfill, P. J., Sell, B. R., Baird, M. A., Allen, J. R., Lavagnino, Z., De Gruiter, H. M., Kremers, G. J., Davidson, M. W., Ustione, A. and Piston, D. W. (2016). Quantitative assessment of fluorescent proteins. Nat. Methods 13, 557–562.

Du, F., Gong, W., Boscá, S., Tucker, M., Vaucheret, H. and Laux, T. (2019). Dose-Dependent AGO1-Mediated Inhibition of the miRNA165/166 Pathway Modulates Stem Cell Maintenance in Arabidopsis Shoot Apical Meristem. Plant Commun. 100002.

Ebhardt, H. A., Fedynak, A. and Fahlman, R. P. (2010). Naturally occurring variations in sequence length creates microRNA isoforms that differ in argonaute effector complex specificity. Silence 1, 1–6.

Fang, X. and Qi, Y. (2015). Rnai in plants: An argonaute-centered view. Plant Cell 28, 272–285.

Gehring, M., Bubb, K. L. and Henikoff, S. (2009). Extensive demethylation of repetitive elements during seed development underlies gene imprinting. Science (80-.). 324, 1447–51.

Gutzat, R., Rembart, K., Nussbaumer, T., Pisupati, R., Hofmann, F., Bradamante, G., Daubel, N., Gaidora, A., Lettner, N., Donà, M., et al. (2018). Stage-specific transcriptomes and DNA methylomes indicate an early and transient loss of transposon control in Arabidopsis shoot stem cells. bioRxiv.

Hamamura, Y., Saito, C., Awai, C., Kurihara, D., Miyawaki, A., Nakagawa, T., Kanaoka, M. M., Sasaki, N., Nakano, A., Berger, F., et al. (2011). Live-cell imaging reveals the dynamics of two sperm cells during double fertilization in Arabidopsis thaliana. Curr. Biol. 21, 497–502.

Havecker, E. R., Wallbridge, L. M., Hardcastle, T. J., Bush, M. S., Kelly, K. A., Dunn, R. M., Schwach, F., Doonan, J. H. and Baulcombe, D. C. (2010). The Arabidopsis RNA-directed DNA methylation argonautes functionally diverge based on their expression and interaction with target loci. Plant Cell 22, 321–34.

He, J., Xu, M., Willmann, M. R., McCormick, K., Hu, T., Yang, L., Starker, C. G., Voytas, D. F., Meyers, B. C. and Poethig, R. S. (2018). Threshold-dependent repression of SPL gene expression by miR156/miR157 controls vegetative phase change in Arabidopsis thaliana. PLoS Genet. 14, 1–28.

Hernández-Lagana, E., Rodríguez-Leal, D., Lúa, J. and Vielle-Calzada, J. P. (2016). A multigenic network of ARGONAUTE4 clade members controls early megaspore formation in arabidopsis. Genetics 204, 1045–1056.

Hsieh, T. T.-F., Ibarra, C. A., Silva, P., Zemach, A., Eshed-Williams, L., Fischer, R. L. and Zilberman, D. (2009). Genome-wide demethylation of Arabidopsis endosperm. Science (80-.). 324, 1451–1454.

Hunter, C., Sun, H. and Poethig, R. S. (2003). The Arabidopsis Heterochronic Gene ZIPPY Is an ARGONAUTE Family Member. Curr. Biol. 13, 1734–1739.

Jouannet, V., Moreno, A. B., Elmayan, T., Vaucheret, H., Crespi, M. D. and Maizel, A. (2012). Cytoplasmic Arabidopsis AGO7 accumulates in membrane-associated siRNA bodies and is required for ta-siRNA biogenesis. EMBO J. 31, 1704–1713.

Jullien, P. E., Susaki, D., Yelagandula, R., Higashiyama, T. and Berger, F. (2012). DNA methylation dynamics during sexual reproduction in Arabidopsis thaliana. Curr. Biol. 22, 1825–1830.

Jullien, P. E., Grob, S., Marchais, A., Pumplin, N., Chevalier, C., Bonnet, D. M., Otto, C., Schott, G. and Voinnet, O. (2020). Functional characterization of Arabidopsis ARGONAUTE 3 in reproductive tissue. Plant J. Accepted Author Manuscript. doi:10.1111/tpj.14868.

Katiyar-Agarwal, S., Gao, S., Vivian-Smith, A. and Jin, H. (2007). A novel class of bacteria-induced small RNAs in Arabidopsis. Genes Dev. 21, 3123–3134.

Kawakatsu, T., Nery, J. R., Castanon, R. and Ecker, J. R. (2017). Dynamic DNA methylation reconfiguration during seed development and germination. Genome Biol. 18, 171.

Landgraf, D., Okumus, B., Chien, P., Baker, T. A. and Paulsson, J. (2012). Segregation of molecules at cell division reveals native protein localization. Nat. Methods 9, 480–482.

Liu, C., Xin, Y., Xu, L., Cai, Z., Xue, Y., Liu, Y., Xie, D., Liu, Y. and Qi, Y. (2018). Arabidopsis ARGONAUTE 1 Binds Chromatin to Promote Gene Transcription in Response to Hormones and Stresses. Dev. Cell 44, 348-361.e7.

Lobbes, D., Rallapalli, G., Schmidt, D. D., Martin, C. and Clarke, J. (2006). SERRATE: a new player on the plant microRNA scene. EMBO Rep. 7, 1052–8.

Lynn, K., Fernandez, A., Aida, M., Sedbrook, J., Tasaka, M., Masson, P. and Barton, M. K. (1999). The PINHEAD/ZWILLE gene acts pleiotropically in Arabidopsis development and has overlapping functions with the ARGONAUTE1 gene. Development 126, 469–481.

Mallory, A. and Vaucheret, H. (2010). Form, function, and regulation of ARGONAUTE proteins. Plant Cell 22, 3879–89.

Marchais, A., Chevalier, C. and Voinnet, O. (2019). Extensive profiling in Arabidopsis reveals abundant polysome-associated 24-nt small RNAs including AGO5-dependent pseudogene-derived siRNAs. Rna 25, 1098–1117.

Martinez, G., Wolff, P., Wang, Z., Moreno-Romero, J., Santos-González, J., Conze, L. L., Defraia, C., Slotkin, R. K. and Köhler, C. (2018). Paternal easiRNAs regulate parental genome dosage in Arabidopsis. Nat. Genet. 50, 193–198.

Matzke, M. A. and Mosher, R. A. (2014). RNA-directed DNA methylation: An epigenetic pathway of increasing complexity. Nat. Rev. Genet. 15, 394–408.

Meinke, D. W. (2020). Genome-wide identification of EMBRYO-DEFECTIVE (EMB) genes required for growth and development in Arabidopsis. New Phytol. 226, 306–325.

Mosher, R. A. and Melnyk, C. W. (2010). siRNAs and DNA methylation: seedy epigenetics. Trends Plant Sci. 15, 204–210.

Moussian, B., Schoof, H., Haecker, A., Ju, G. and Laux, T. (1998). Role of the ZWILLE gene in the regulation of central shoot meristem cell fate during Arabidopsis embryogenesis. Curr. Opin. Plant Biol. 1, 188.

Nodine, M. D. and Bartel, D. P. (2010). MicroRNAs prevent precocious gene expression and enable pattern formation during plant embryogenesis. Genes Dev. 24, 2678–92.

Oliver, C., Santos, J. L. and Pradillo, M. (2014). On the role of some ARGONAUTE proteins in meiosis and DNA repair in Arabidopsis thaliana. Front. Plant Sci. 5, 177.

Olmedo-Monfil, V., Durán-Figueroa, N., Arteaga-Vázquez, M., Demesa-Arévalo, E., Autran, D., Grimanelli, D., Slotkin, R. K., Martienssen, R. A. and Vielle-Calzada, J.-P. (2010). Control of female gamete formation by a small RNA pathway in Arabidopsis. Nature 464, 628–632.

Rodríguez-Leal, D., León-Martínez, G., Abad-Vivero, U. and Vielle-Calzada, J. P. (2015). Natural variation in epigenetic pathways affects the specification of female gamete precursors in arabidopsis. Plant Cell 27, 1034–1045.

Roussin-Léveillée, C., Silva-Martins, G. and Moffett, P. (2020). ARGONAUTE5 Represses Age-Dependent Induction of Flowering through Physical and Functional Interaction with miR156 in Arabidopsis. Plant Cell Physiol. 61, 957–966.

Schröder, J. A. and Jullien, P. E. (2019). The Diversity of Plant Small RNAs Silencing Mechanisms. Chim. Int. J. Chem. 73, 362–367.

Seefried, W. F., Willmann, M. R., Clausen, R. L. and Jenik, P. D. (2014). Global regulation of embryonic patterning in arabidopsis by MicroRNAs. Plant Physiol. 165, 670–687.

Sharma, U., Sun, F., Conine, C. C., Reichholf, B., Kukreja, S., Herzog, V. A., Ameres, S. L. and Rando, O. J. (2018). Small RNAs Are Trafficked from the Epididymis to Developing Mammalian Sperm. Dev. Cell 46, 481-494.e6.

Slotkin, R. K., Vaughn, M., Borges, F., Tanurdzic, M., Becker, J. D. J. D., Feijo, J. A., Martienssen, R. A., Tanurdžić, M., Feijó, J. A., Feijo, A., et al. (2009). Epigenetic reprogramming and small RNA silencing of transposable elements in pollen. Cell 136, 461–472.

Sprunck, S., Urban, M., Strieder, N., Lindemeier, M., Bleckmann, A., Evers, M., Hackenberg, T., Möhle, C., Dresselhaus, T. and Engelmann, J. C. (2019). Elucidating small RNA pathways in Arabidopsis thaliana egg cells. bioRxiv.

Stroud, H., Greenberg, M. V. C., Feng, S., Bernatavichute, Y. V. and Jacobsen, S. E. (2012). Resource Comprehensive Analysis of Silencing Mutants Reveals Complex Regulation of the Arabidopsis Methylome. Cell 152, 352–364.

Takeda, A., Iwasaki, S., Watanabe, T., Utsumi, M. and Watanabe, Y. (2008). The mechanism selecting the guide strand from small RNA duplexes is different among Argonaute proteins. Plant Cell Physiol. 49, 493–500.

Tucker, M. R., Hinze, A., Tucker, E. J., Takada, S., Jürgens, G. and Laux, T. (2008). Vascular signalling mediated by ZWILLE potentiates WUSCHEL function during shoot meristem stem cell development in the Arabidopsis embryo. Development 135, 2839–2843.

Tucker, M. R., Okada, T., Hu, Y., Scholefield, A., Taylor, J. M. and Koltunow, A. M. G. (2012). Somatic small RNA pathways promote the mitotic events of megagametogenesis during female reproductive development in Arabidopsis. Development 139, 1399–1404.

Van Ex, F., Jacob, Y. and Martienssen, R. a (2011). Multiple roles for small RNAs during plant reproduction. Curr. Opin. Plant Biol. 1–6.

Vazquez, F., Gasciolli, V., Crété, P. and Vaucheret, H. (2004). The Nuclear dsRNA Binding Protein HYL1 Is Required for MicroRNA Accumulation and Plant Development, but Not Posttranscriptional Transgene Silencing. Curr. Biol. 14, 346–351.

Weick, E. M. and Miska, E. A. (2014). piRNAs: From biogenesis to function. Dev. 141, 3458–3471.

Willmann, M. R., Mehalick, A. J., Packer, R. L. and Jenik, P. D. (2011). MicroRNAs regulate the timing of embryo maturation in Arabidopsis. Plant Physiol. 155, 1871–1884.

Wu, G., Park, M. Y., Conway, S. R., Wang, J. W., Weigel, D. and Poethig, R. S. (2009). The Sequential Action of miR156 and miR172 Regulates Developmental Timing in Arabidopsis. Cell 138, 750–759.

Wuest, S. E., Vijverberg, K., Schmidt, A., Weiss, M., Gheyselinck, J., Lohr, M., Wellmer, F., Rahnenführer, J., von Mering, C. and Grossniklaus, U. (2010). Arabidopsis Female Gametophyte Gene Expression Map Reveals Similarities between Plant and Animal Gametes. Curr. Biol. 20, 506–512.

Ye, R., Wang, W., Iki, T., Liu, C., Wu, Y., Ishikawa, M., Zhou, X. and Qi, Y. (2012). Cytoplasmic Assembly and Selective Nuclear Import of Arabidopsis ARGONAUTE4/siRNA Complexes. Mol. Cell 46, 859–870.

Yuan, S., Schuster, A., Tang, C., Yu, T., Ortogero, N., Bao, J., Zheng, H. and Yan, W. (2016). Sperm-borne miRNAs and endo-siRNAs are important for fertilization and preimplantation embryonic development. Dev. 143, 635–647.

Zhang, H., Xia, R., Meyers, B. C. and Walbot, V. (2015). Evolution, functions, and mysteries of plant ARGONAUTE proteins. Curr. Opin. Plant Biol. 27, 84–90.

Zhao, L., Cai, H., Su, Z., Wang, L., Huang, X., Zhang, M., Chen, P., Dai, X., Zhao, H., Palanivelu, R., et al. (2018a). KLU suppresses megasporocyte cell fate through SWR1-mediated activation of WRKY28 expression in Arabidopsis. Proc. Natl. Acad. Sci. U. S. A. 115, E526–E535.

Zhao, Y., Wang, S., Wu, W., Li, L., Jiang, T. and Zheng, B. (2018b). Clearance of maternal barriers by paternal miR159 to initiate endosperm nuclear division in Arabidopsis. Nat. Commun. 9, 5011.

Zheng, X., Zhu, J., Kapoor, A. and Zhu, J.-K. (2007). Role of Arabidopsis AGO6 in siRNA accumulation, DNA methylation and transcriptional gene silencing. EMBO J. 26, 1691–1701.

